# Distinct features of the *Leishmania* cap-binding protein LeishIF4E2 revealed by CRISPR-Cas9 mediated heterozygous deletion

**DOI:** 10.1101/2020.05.13.093914

**Authors:** Nofar Baron, Nitin Tupperwar, Geula Davidov, Raz Zarivach, Michal Shapira

**Author notes:** Corresponding Author: Prof. Michal Shapira, Ph.D., Department of Life Sciences, Ben-Gurion University of the Negev, Beer Sheva, Israel. Authors contributed equally (Nofar Baron, Nitin Tupperwar).

## Abstract

*Leishmania* parasites cycle between sand-fly vectors and mammalian hosts, adapting to alternating environments by stage-differentiation, accompanied by changes in the proteome profiles. Translation regulation plays a central role in driving the differential program of gene expression, since control of gene regulation in *Leishmania* is mostly post-transcriptional. The *Leishmania* genome encodes six eIF4E candidates, each assumed to have specific functions, although overlaps are expected. It is noted that some of them can bind to a dedicated eIF4G candidate partners, and LeishIF4E2 does not bind any known eIF4G ortholog. LeishIF4E2 was previously shown to comigrate in the polysomal fractions of sucrose gradients, whereas initiation factors usually comigrate with pre-initiation and initiation complexes. Using the CRISPR-Cas9 methodology, we deleted one of the two LeishIF4E2 gene copies. The deletion caused severe alterations in the morphology of mutant cells that turned round and equipped with a very short flagellum, but their growth rate and general translation remained unaffected. Proteomic analysis of the LeishIF4E2(+/-) mutant cells compared to wild type controls showed that the number of proteins that were upregulated exceeded the number of downregulated proteins, possibly indicating that a repressor function was eliminated. The upregulated proteins were related mainly to general metabolic processes, DNA repair and replication, signaling, and cellular motor activity. The downregulated proteins included several groups, including cytoskeletal and ribosomal proteins. Despite the fact that only one of the two LeishIF4E2 alleles was deleted, the mutant cells were impaired in their ability to infect cultured macrophages. LeishIF4E2 does not behave like a general translation factor and its function remains elusive. Although it could have a repressive function, we cannot exclude the possibility that it is responsible for translation of a specific set of transcripts. Overall, our results are in line with the possibility that the different LeishIF4Es are assigned unique functions.

**Author summary:** *Leishmania* parasites cause a broad spectrum of diseases that lead to different pathological symptoms. During their life cycle, the parasites shuffle between sand-fly vectors and mammalian hosts, while adapting to changing environments via a stage specific program of gene expression that assists the survival of *Leishmania* under the changing conditions. Translation initiation plays a key role in control of gene expression, in *Leishmania* this is exemplified by the presence of multiple cap-binding complexes that interact with mRNAs. The parasites encode multiple paralogs of the cap-binding translation initiation factor eIF4E, and of its corresponding binding partner eIF4G, forming complexes with different potential functions. Using the CRISPR-Cas9 methodology, we generated a heterozygous mutant of the least studied cap-binding paralog, LeishIF4E2, eliminating one of its two alleles. The mutation caused morphological defects, resulting in short and rounded up cells with a significant reduction in their flagellar length. Although general translation rates and growth of the mutant parasite were not affected, total proteome analysis and translation assay suggested that LeishIF4E2 could possibly function as a translation repressor. Alternatively, LeishIF4E-2 could be responsible for promoting translation of a specific set of transcripts.

## Introduction

*Leishmania* species are unicellular trypanosomatid protists that run a digenetic life cycle, migrating between sand flies and mammals [1]. In the sand fly, the parasites reside in the alimentary canal, as extra-cellular promastigotes, attached to the gut wall. Promastigotes are elongated and equipped with a flagellum that enables them to attach to the gut wall, and further promotes their mobility towards the front parts of the mouth during metacyclogenesis [1]. Metacyclic promastigotes are transmitted into the mammalian host during the females’ blood meal, where they enter into macrophages and differentiate into the non-flagellated amastigotes, residing within phagolysosomal vacuoles. Amastigotes are smaller in size, rounded up and their flagellum does not protrude from the flagellar pocket. Amastigotes are therefore non-motile. During their life cycle, *Leishmania* parasites are exposed to non-favorable environments and extreme conditions [2, 3].

Translation in eukaryotes proceeds mostly by cap-dependent mechanisms which are primarily controlled at the level of initiation [4]. The eIF4F complex binds to the mRNA that consists of the cap-binding protein (eIF4E), an RNA-helicase (eIF4A) and a scaffold protein (eIF4G) that serves as a hub that holds together the pre-initiation complex (PIC). Assembly of the cap-binding complex is globally controlled by 4E-binding proteins, such as 4E-BP, a small (10 KDa) and unstructured protein, which contains the consensus Y(X_4_)LΦ motif for binding to eIF4E. 4E-BP competes with eIF4G for binding to eIF4E therefore preventing assembly of the translation initiation complex [5]. Other 4E-binding proteins can control the cap-dependent translation either globally, or in a transcript-specific manner, when an inhibitory complex that assembles over the cap-binding protein is stabilized through proteins that interact with elements in the 3’ UTR [6, 7]. Another mode of regulation involves an ortholog of eIF4E, denoted 4E-HP, which binds to the cap-structure, but not to any eIF4G, competing with the canonical eIF4E to bind to the mRNA cap-structure [8].

There are six paralogs of the cap-binding protein eIF4E in *Leishmania*, designated LeishIF4Es 1 to 6. They vary in their cap-binding affinities and were reported to have diverse functions [9, 10]. Three paralogs of eIF4A were reported in *Leishmania*, [11] and five eIF4G candidates containing the MIF4G domain (known as the middle domain of eIF4G) have been identified [9, 11, 12].

LeishIF4E4 was suggested to be a canonical cap-binding protein in promastigotes form. LeishIF4E4 is a cap-binding protein that forms a complex with LeishIF4G3 and LeishIF4A1. LeishIF4E4 also binds directly to LeishPABP1 through its extended N-terminus. This interaction was recently confirmed by the solved 3-D structure describing this interaction [13]. LeishIF4E1 is a strong cap-binding protein, but does not bind to any of the known LeishIF4G isoforms. It is the only cap binding protein that maintains its expression and binding affinity to m^7^GTP in axenic amastigotes [14]. Although LeishIF4E1 does not form a complex with any of the LeishIF4G candidates, a recent report based on CRISPR-Cas9 deletion, suggested that despite the absence of a functional eIF4G binding partner, LeishIF4E1 appears to promote translation [15]. The 3-D structure of LeishIF4E1 was recently solved, when it was associated with a fragment derived from its unique binding partner Leish4E-IP1. Semi-*in vivo* binding assays suggested that recombinant Leish4E-IP1 inhibits the cap-binding activity of LeishIF4E1 [16], suggesting that 4E-IP1 is a functional inhibitor of LeishIF4E1.

LeishIF4E3 has a weak cap-binding activity and is known to bind LeishIF4G4, one of the MIF4G proteins in *Leishmania* and *Trypanosoma* [10, 17, 18]. However, upon nutritional stress that mimics metacyclogenesis, LeishIF4E3 was found to concentrate in cytoplasmic granules. These granules function as storage sites for inactive mRNAs and ribosomal proteins [19]. Furthermore, deletion of one copy of LeishIF4E3 by the CRISPR-Cas9 system changed the cell morphology and reduced the infectivity of *Leishmania* cells, while the attempt to generate null mutant did not succeed [20].

Two additional cap-binding proteins were identified in *T. brucei*. TbIF4E5 interacts with TbIF4G1 and TbIF4G2 and has a role in the motility of parasite. TbIF4E5 associates with a different set of the complex with either TbIF4G1 and TbIF4G2, but the role of each of the complex in terms of translation is not clear [21]. TbEIF4E6 forms a tripartite cytosolic complex with TbEIF4G5 and with a 70.3 kDa hypothetical protein, TbG5-IP, that contains domains that are typical of two capping enzymes, the nucleoside triphosphate hydrolase and guanylyltransferase. Although the exact role of this complex is not known, it could be involved in recovery of decapped mRNAs [22].

LeishIF4E2 is the least studied paralog in *Leishmania*. It contains the three conserved tryptophan residues that are part of the cap-binding pocket. Binding assays based on recombinant proteins showed that LeishIF4E2 binds better to cap-4 and less to m^7^GTP, explaining why it could not be affinity-purified efficiently over m^7^GTP-Sepharose [10]. LeishIF4E2 co-migrated with high molecular weight fractions on sucrose gradients. Treatment of the cell extracts with mung-bean nuclease caused the protein to migrate at the top of the gradient, suggesting that it was most probably associated with polysomes [10]. Previous analyses of the canonical translation initiation factors in eukaryotes showed that the preinitiation complexes concentrated mostly in the 43S fractions over sucrose gradients [23], yet LeishIF4E2 was rather different in that sense as it migrated with the heavy fractions.

The *T. brucei* ortholog of LeishIF4E2, TbEIF4E2, also fails to bind to any TbEIF4G homolog [24]. However, it was reported to bind mature mRNAs and to associate with the histone mRNA stem-loop binding protein (SLBP). Two SLBPs were found in *T. brucei*, SLBP1 and SLBP2, the latter is unique to trypanosomes. SLBP2 binds TbEIF4E2 through an exclusive region that was found in SLBP2. SLBP2 and TbEIF4E2 are cytoplasmic proteins, which are more abundant in early-log phase [24]. Despite the available information about LeishIF4E2, the ortholog of TbEIF4E2, its role in translation is vague. The development of the CRISPR-Cas9 mediated knock-out system for *Leishmania* allowed us to easily proceed towards targeted gene knock-outs, either complete, or partial.

Using the CRISPR-Cas9 system for *Leishmania*, we generated a heterozygous mutant of LeishIF4E2, in which a single allele was deleted. Global translation and growth of the LeishIF4E2(+/-) mutant cells were not affected, but normal parasite morphology was altered, as the mutant cells almost lost their flagella leaving a short rudiment, and became round in shape. The mass spectrometry analysis monitoring the total proteome of the LeishIF4E2 mutant indicated that a larger number of proteins were upregulated as compared to controls. Finally, we also report that LeishIF4E2 expression is required for parasite infectivity as measured by their entry and sustainability within macrophages, as infectivity of LeishIF4E2(+/-) mutant was reduced.

## Materials and methods

### Cells

*Leishmania mexicana* M379 cells were routinely cultured at 25°C in Medium 199 (M199), pH 7.4, supplemented with 10% fetal calf serum (FCS, Biological Industries), 5 µg/mL hemin, 0.1 mM adenine, 40 mM Hepes, 4 mM L-glutamine, 100 U/mL penicillin and 100 µg/mL and streptomycin.

RAW 264.7 macrophage cells were grown at 37°C in DMEM supplemented with 10% FCS, 4 mM L-glutamine, 0.1 mM adenine, 40 mM Hepes pH 7.4, 100 U/mL penicillin and 100 µg/mL streptomycin, in an atmosphere of 5% CO2.

### Affinity purification of recombinant LeishIF4E2

LeishIF4E2 was cloned in the pET28 vector and expressed in Rossetta strain of *E. coli.* The bacterial cells were grown to the density of OD600 =0.5–0.7 and expression was induced by the addition of 0.3 mM IPTG at 25°C for 8 hrs. The cells were harvested, and re-suspended in lysis buffer [20 mM HEPES-KOH pH 7.5, 100 mM KCl, 2 mM Tris (2-carboxyethyl) phosphine hydrochloride (TCEP), 0.1 EDTA, 0.01% Triton X-100, protease inhibitors and 5 µg/mL DNaseI. Ni-NTA beads (Cube Biotech) (5 mL) were pre-washed with Binding Buffer [20 mM HEPES-KOH pH 7.5, 150 mM KCl, 2 mM TCEP; 2 column volumes (CV)]. The cells were disrupted using a French Press at 1500 psi, followed by centrifugation at 45,000 rpm (Beckman 70 Ti rotor). The supernatant was further incubated with Ni-NTA beads. The first wash was carried out with Wash Buffer I (WB I, 20 mM Hepes-KOH pH 7.5, 750 mM KCl, 2 mM TCEP and 10 mM imidazole, in 2 CVs). The second wash was performed with WB II (20 mM Hepes-KOH pH 7.5, 500 mM KCl, 2 mM TCEP and 10 mM imidazole, in 2 CVs). The third wash was done with WB III (20 mM Hepes-KOH pH 7.5, 250 mM KCl, 2 mM TCEP and 20 mM imidazole 1 CV). The fourth wash was performed with WB IV (20 mM Hepes-KOH pH 7.5, 100 mM KCl, 2 mM TCEP and 30 mM imidazole, 1 CV). Finally, elution was done by passing one volume of 20 mM Hepes-KOH pH 7.5, 100 mM KCl, 2 mM TCEP and 300 mM imidazole. The Recombinant protein was dialyzed overnight at 4°C against the Binding Buffer.

### CRISPR-Cas9 mediated deletion of single allele of LeishIF4E2

Plasmids developed for the CRISPR system in *Leishmania* were obtained from Dr. Eva Gluenz [University of Oxford, UK, [25]]. The pTB007 plasmid contained the genes encoding the *Streptococcus pyogens* CRISPR-associated protein 9 endonuclease and the T7 RNA-polymerase gene (Cas9/T7), along with the hygromycin resistance gene, between upstream and downstream sequences that generated its UTRs. pTB007 was transfected into *L. mexicana* promastigotes and transgenic cells stably expressing the Cas9 and T7 RNA-polymerase were selected for hygromycin resistance (200 μg/mL).

### Generation of LeishIF4E2(+/-) mutant by CRISPR-Cas9

To generate LeishIF4E2(+/-) mutants, we used three PCR amplified products. These were the two 5’ and 3’ sgRNAs designed to create double-strand breaks upstream and downstream of the LeishIF4E2 coding region, and the LeishIF4E2 repair cassette fragment, containing the G418 resistance marker. The three PCR products (4 µg of each) were transfected into mid-log phase transgenic cells expressing Cas9 and T7 RNA-polymerase, and cells were further selected for resistance to 200 μg/mL G418 [26].

The sgRNA sequences that were used to delete the LeishIF4E2 gene were obtained from LeishGEdit.net [27]. The sgRNAs contained the highest-scoring 20nt sequence within 105 bp upstream or downstream of the target gene. The sequences of the sgRNAs were blasted against the *L. mexicana* genome in TriTrypDB, to verify that the sgRNAs were exclusively specific for LeishIF4E2(E value = 0.001 and 8e^-5^). We also ran a BLAST analysis with the drug resistance repair cassette that contained the homology sequence to the UTR of LeishIF4E2, to specifically target the insertion of the selection marker. The repair cassette showed an E value of 5e^-9^, suggesting very high specificity of the system. The sgRNA target sequences and the homology arms on the repair cassette fully matched the target sequence of LeishIF4E2.

### PCR amplification of sgRNA templates

DNA fragments encoding LeishIF4E2 specific 5’ and 3’ guide RNAs for cleavage upstream and downstream to the LeishIF4E2 target gene were generated. The template for this PCR reaction consisted of two fragments, one contained the common sgRNA scaffold fragment (5’AAAGCACCGACTCGGTGCCACTTTTTCAAGTTGATAACGGACTAGCCTTATTT TAACTTGCTATTTCTAGCTCTAAAAC-3’) and the other contained the T7 RNA polymerase promoter (small letters) fused to the gRNA (5’ or 3’), targeting LeishIF4E2(capital letters) and a short sequence overlapping with the scaffold fragment (small letters). Thus, the two individual template fragments for targeting a double-strand break of LeishIF4E2 were (5’-gaaattaatacgactcactataggGTGAAGCGCGTTTCATTTCCgttttagagctagaaatagc-3’) for the 5’ cleavage site and (5’-gaaattaatacgactcactataggACGCTGAGGTTGACAGGTCGgttttagagctagaaatagc-3’) for the 3’ cleavage site. Each of these two fragments (1 µM each) was annealed to the partially overlapping scaffold fragment and further amplified with two small primers (2 µM each), derived from the T7 promoter (G00F, 5’-TTAATACGACTCACTATAGG-3’) and the common scaffold fragment (G00R, 5’-GCACCGACTCGGTGCCACTT-3’). The reaction mixture consisted of dNTPs (0.2 mM), HiFi Polymerase (1 unit, Phusion, NEB) in GC buffer with MgCl_2_ (NEB), in a total volume of 50 µL. The PCR conditions included an initial denaturation at 98°C for 2 min, followed by 35 cycles of 98°C for 10 s, annealing at 60°C for 30 s and extension at 72°C for 15 s. All PCR products were gel-purified and heated at 94°C for 5 minutes before transfection.

### PCR amplification of the LeishIF4E2 replacement fragment

A DNA fragment designed to repair the double-strand breaks surrounding the LeishIF4E2 target gene was amplified by PCR. The LeishIF4E2 specific primers were derived from the 5’ and 3’ endogenous UTR sequences upstream and downstream to the LeishIF4E2 gene and the sequences from the antibiotic repair cassette, based on the LeishGEdit database (http://www.leishgedit.net/Home.html). The primers were (5’-ATAGACATCGCATCAGGTTTTTTTTGCTTCgtataatgcagacctgctgc -3’, forward) and 5’-GCAGATAAGGGAATATCTGACAGGCGGCGCccaatttgagagacctgtgc-3’, reverse). Capital letters represent the UTR sequences of the LeishIF4E2 gene and small letters represent the region on the pT plasmid that flank the UTR adjacent to the antibiotic resistance gene. The PCR for generating the fragment used for repair of the double-strand breaks on both sides of the gene targeted for deletion, was performed using the pTNeo plasmid as a template. The resulting fragments promote the integration of the drug resistance marker by homologous recombination at the target site. The reaction mixture consists of 2 µM of each primer, dNTPs (0.2 mM), the template pTNeo (30 ng), 3% (v/v) DMSO, HiFi Polymerase (1 unit of Phusion, NEB) in GC buffer (containing MgCl_2,_ to a final concentration of 1.5mM) in a total volume of 50 µL. PCR conditions included initial denaturation at 98°C for 4 min which is followed by 40 cycles of 98°C for 30 Seconds (s), annealing at 65°C for 30 s and extension at 72°C for 2 min 15 s. The final extension was performed during 7 min at 72°C. All PCR products were gel purified and heated at 94°C for 5 minutes before transfection.

### Diagnostic PCR to confirm the deletion of LeishIF4E2

Genomic DNA from the drug-resistant cells was isolated 14 days post transfection using DNeasy Blood & Tissue Kit (Qiagen) and analyzed for the presence of the LeishIF4E2 gene, using specific primers derived from the untranslated regions (UTR) of the LeishIF4E2. The primers used were LeishIF4E2 (5’UTR) forward (5’-CGTAGCTCAGATAGACTA-3’) and LeishIF4E2 reverse (3’UTR) (5’-AGACATCAGAGTATTCGT -3’). A parallel reaction was performed to look for the presence of the G418 resistance gene, with primers derived from its ORF: G418 F (5’-GCCCGGTTCTTTTTGTCAAGAC-3’) and G418 R (5’-GTCACGACGAGATCATCATCGCCG-3’). Genomic DNA from Cas9/T7 *L. mexicana* cells was used to detect the presence of the LeishIF4E2 gene. The reaction mixture consisted of 2 µM of each primer, gDNA (100 ng), dNTPs (0.2 mM), HiFi Polymerase (1 unit, Phusion, NEB) in GC buffer with MgCl_2_ (NEB) in a total volume of 50 µl. PCR conditions included initial denaturation at 98°C for 4 min which is followed by 35 cycles of 98°C for 30 s, annealing at 60°C for 30 s and extension at 72°C for 2 min 15 s. Final extension was done for 7 min at 72°C. PCR products were separated over 1% agarose gels.

### Generation of LeishIF4E2 add-back parasites

The transgenic LeishIF4E2(+/-) deletion mutant cells were transfected with an episomal transfection vector that promoted the recovery of LeishIF4E2 expression. The plasmid was derived from pTPuro, which confers resistance to puromycin. The LeishIF4E2 added-back gene from *L. mexicana* was tagged with the Streptavidin binding peptide (SBP, ∼4 kDa), enabling its further identification in the transgenic parasites by antibodies against the SBP tag [14]. The tagged LeishIF4E2 was cloned between two intergenic regions derived from the HSP83 (H) genomic cluster. The pTPuro-LeishIF4E2 plasmid was generated as follows: The ORF of LeishIF4E2 from *L. mexicana* was amplified using the forward 5’-GGATCCATGGACCCGAGTAGGTGTG -3’ and reverse 5’-TCTAGA CAC AAC CTC ACC CCT CAC A-3’ primers, with BamHI and XbaI sites introduced at the 5’ ends of these primers (small letters). The BamHI/XbaI PCR product of LeishIF4E2 was cloned into the BamHI and XbaI sites of the pTPuro that already contained LeishIF4E1-SBP between two intergenic regions derived from the HSP83 genomic locus [28, 29]. The resulting pTPuro-LeishIF4E2-SBP expression vector was transfected into the LeishIF4E2(+/-) deletion mutant, and cells were selected for their resistance to Puromycin (100 µg/mL).

### Growth analysis

*L. mexicana* M379 wild type and Cas9/T7 expressing cells, along with the LeishIF4E2(+/-) deletion mutant were cultured as promastigotes at 25°C in M199 medium containing all supplements **(see above).** Cells were seeded at a concentration of 5×10^5^ cells/mL, and counted daily during 5 consecutive days. The curves were obtained from three independent repeats.

### Western analysis

Cells at their mid-log phase of growth (10 mL) were harvested, washed twice with phosphate buffered saline (PBS) and once in post ribosomal supernatant [(PRS) buffer, 35 mM Hepes pH 7.5, 100 mM KCl, 10 mM MgCl2, 1 mM DTT]. The cell pellet was resuspended in PRS+ (300 ul), which was PRS supplemented with a 2x cocktail of protease inhibitors (Sigma) and 4 mM iodoacetamide (Sigma), along with (+) phosphatase inhibitors: 25 mM sodium fluoride, 55 mM β-glycerophosphate and 5 mM sodium orthovanadate. Cells were lysed by the addition of 65 µl of X5 Laemmli sample buffer and heated at 95°C for 5 minutes. Cell extracts (40 uL) were resolved over 10% SDS-PAGE, blotted and further probed using specific primary and secondary antibodies.

Antibodies against LeishIF4E2(rabbit polyclonal, 1:2,000) and against the SBP tag (Millipore, monoclonal, 1:10,000), were used to detect the endogenous and tagged LeishIF4E2 proteins, respectively. The blots were developed incubation with specific peroxidase-labelled secondary antibodies against rabbit (KPL, 1:10,000 for LeishIF4E2) and mouse (KPL, 1:10,000 for SBP).

### Translation assay

Global translation levels were monitored using the non-radioactive SUnSET (Surface SEnsing of Translation) assay. This assay is based on the incorporation of puromycin, an amino-acyl tRNA analog, into the A site of translating ribosomes [30]. Cells were incubated with puromycin (1 µg/mL, Sigma) during 30 min, then washed twice with PBS and once with PRS+. Cell pellets were resuspended in 300 µl of PRS^+^ buffer, denatured in Laemmli sample buffer and boiled for 5 min. Cells treated with cycloheximide (100 µg/mL) prior to the addition of puromycin served as a negative control. Samples were resolved over 10% SDS-Polyacrylamide gel electrophoresis (SDS-PAGE). The gels were blotted and subjected to western analysis using monoclonal mouse anti-puromycin antibodies (DSHB, 1:1,000) and secondary peroxidase labeled anti-mouse antibodies (KPL, 1:10,000).

### Phase contrast microscopy of *Leishmania* promastigotes

Cells from different lines in their late-log phase of growth were harvested, washed in cold PBS, fixed in 2% paraformaldehyde in PBS and mounted onto glass slides. Phase contrast microscope images were captured at X100 magnification with a Zeiss Axiovert 200M microscope equipped with AxioCam HRm CCD camera.

### Flow cytometry analysis of *Leishmania*

Cell viability was verified by incubation of the cells with 20 µg/mL propidium iodide (PI) during 30 minutes. The stained cells were analyzed using the ImageStream X Mark II Imaging flow cytometer (Millipore) with a X60/0.9 objective. Data from channels representing bright field, and fluorescence (PI) emission at 488 nm (to evaluate cell viability) were recorded for 20,000 cells for each analyzed sample. IDEAS software generated the quantitative measurements of the focused, single and live cells for all the four examined cell strains. Cell shape was quantified using circularity and elongatedness features applied on the bright field image processed by adaptive erode mask. Representative scatter plots are shown for focused single cells, and for circularity (cell shape). Recorded emission of the PI in the gated population evaluated cell viability.

### Data analysis

IDEAS software was used to generate the quantitative measurements of images recorded for the examined cell population. The focus quality of each cell was first determined by measuring the gradient root mean square (RMS) value. The cells representing high RMS value in the histogram were gated, to select cells in focus. In the second step, single cell populations were gated from the scatter plot of aspect ratio/area, to exclude cell aggregates. Further, the intensity of PI staining was used to exclude dead cells. The remaining living, single cells in focus were subjected to image analysis to determine cell morphology. To obtain cell shape, a customized adaptive erode mask was used on the bright field channel, with a coefficient of 78. We further customized this mask to exclude the flagellum from the cell shape analysis. Further, circularity and elongatedness features were measured. A predetermined threshold value of 4 was set to define circularity. Elongatedness values represent the ratio between cell length and width. Representative scatter plots are presented for focused single cells, and for circularity. Cell viability was measured by recording the emission of PI in the gated population. All data shown are from a minimum of three biological replicates. A similar approach was taken for generating a template adapted for measuring the structural features of axenic amastigotes. Furthermore, due to the tendency of axenic amastigotes to aggregate, different cell populations were gated to obtain only the single rounded cells (resembling axenic amastigotes). The gated area excluded cell aggregates, debris and elongated promastigotes in a specific area of scatter plot. This template was further used to quantify the single round amastigote like cells in all the cell lines that were analyzed.

### *In vitro* macrophage infection assay

*L. mexicana* LeishIF4E2(+/-) mutants and add-back cells, wild type and transgenic parasites expressing Cas9/T7 polymerase, were seeded at the concentration of 5×10^5^ cells/ mL and allowed to grow for 5 days, to reach their stationary growth phase. Wild type, Cas9/T7 expressers, LeishIF4E2(+/-) and add-back were washed with DMEM (Dulbecco’s Modified Eagle Medium) and labeled by incubation with 10 μM carboxyfluorescein succinimidyl ester (CFSE) in DMEM at 25°C for 10 mins. The cells were then washed with DMEM, counted and used to infect RAW 264.7 macrophages. The macrophages (5×10^5^) were pre-seeded a day in advance in chambered slides (Ibidi)and incubated with the parasites at a ratio of 10:1 during 1 hour, in 300 µl. The cells were then washed three times with PBS and once in DMEM, to remove extracellular parasites. The infected macrophages were either fixed immediately for further analysis by confocal microscopy, or further incubated for 24h at 37°C in an atmosphere containing 5% CO_2_. The infected macrophages were then processed for confocal microscopy as described below. A single representative section of Z-projections (maximum intensity) produced by Image J software is presented in all the figures. The infectivity values were determined using the cell counting plugin in ImageJ. We first counted the number of infected cells in a total of 200 macrophages, and then counted the number of internalized parasites within the infected cells, 1 or 24 hours following infection. Statistical analysis was generated using GraphPad Prism 5. We used the non-parametric Kruskal-Wallis test to determine significant differences in the infectivity and in the average number of parasites per infected macrophages.

### Confocal microscopy of *Leishmania* promastigotes

Infected macrophages, following 1 or 24 hrs of infection, were washed with PBS, fixed in 2% paraformaldehyde for 30 min, washed once with PBS and permeabilized with 0.1% Triton X-100 in PBS, for 10 min. Nucleic acids were stained with 4′,6-diamidino-2-phenylindole (DAPI, Sigma, 1 µg/mL) and the cells were washed three times with PBS. The slides were observed using an inverted Zeiss LSM 880 Axio-observer Z1 confocal laser scanning microscope with Airyscan detector. Cells were visualized using Zeiss Plan-Apochromat oil objective lens of X63 and numerical aperture of 1.4. Z-stacked images were acquired with a digital zoom of X8 (X1.8 for broad fields), using the Zen lite software (Carl Zeiss microscopy). Images were processed using Image J software package. A single representative section of the compiled Z-projections produced by Image J software is presented in all the figures.

### Mass spectrometry analysis

To characterize the proteomic differences between the LeishIF4E2(+/-) deletion mutant and wild type cells, we performed a mass spectrometry analysis of total cell lysates prepared from the LeishIF4E2(+/-) mutant cells. Total cell lysates from mid log stage promastigotes WT, Cas9/T7, LeishIF4E2(+/-) were resuspended in a buffer containing 100 mM Tris HCl pH 7.4, 10 mM DTT, 5% SDS, 2 mM iodoacetamide and a cocktail of protease inhibitors (Sigma). Cell lysates were precipitated using 10 % trichloro-acetic acid (TCA) and the pellets were washed with 80% acetone. The mass spectrometric analysis was performed by the Smoler Proteomics Center at the Technion, Israel.

#### Mass spectrometry (MS)

Proteins were reduced using 3 mM DTT (60°C for 30 min), followed by modification with 10 mM iodoacetamide in 100 mM ammonium bicarbonate for 30 min at room temperature. This was followed by overnight digestion in 10 mM ammonium bicarbonate in trypsin (Promega) at 37°C. Trypsin-digested peptides were desalted, dried, resuspended in 0.1 % formic acid and resolved by reverse phase chromatography over a 30 min linear gradient with 5% to 35% acetonitrile and 0.1 % formic acid in water, a 15 min gradient with 35% to 95% acetonitrile and 0.1 % formic acid in water and a 15 min gradient at 95% acetonitrile and 0.1 % formic acid in water at a flow rate of 0.15 µl/min. MS was performed using a Q-Exactive Plus Mass Spectrometer (Thermo) in positive mode set to conduct a repetitively full MS scan followed by high energy collision dissociation of the 10 dominant ions selected from the first MS scan. A mass tolerance of 10 ppm for precursor masses and 20 ppm for fragment ions was set.

#### Statistical analysis for enriched proteins

Raw mass spectrometric data were analyzed by the MaxQuant software, version 1.5.2.8 [31]. The data were searched against the annotated *L. mexicana* proteins from the TriTrypDB [32]. Protein identification was set at less than a 1% false discovery rate. The MaxQuant settings selected were a minimum of 1 razor/unique peptide for identification, with a minimum peptide length of six amino acids and a maximum of two mis-cleavages. For protein quantification, summed peptide intensities were used. Missing intensities from the analyses were substituted with values close to baseline only if the values were present in the corresponding analyzed sample. The log_2_ of LFQ intensities [33] were compared between the three biological repeats of each group on the Perseus software platform [34], using a t-test. The enrichment threshold was set to a log_2_ fold change > 0.8 and p < 0.05. The annotated proteins were first categorized manually.

#### Categorization of enriched proteins by the gene ontology (GO) annotation via TriTrypDB

Enriched proteins were classified by the GO Annotation tool in TriTrypDB, based on molecular functions. The threshold for the calculated enrichment of proteins based on their GO terms was set at 2.5 fold, with a p<0.05. This threshold eliminated most of the general groups that represented parental GO terms. GO terms for which only a single protein was annotated were filtered out as well. In some cases, GO terms that were included in other functional terms are not shown, leaving only the representative GO term

### Statistical analysis

Statistical analysis was performed using GraphPad Prism version 5. Each experiment was performed independently at least three times and the individual values were presented as dots. For experiments with a higher number of repeats, results are expressed as Mean ± SD. Statistical significance was determined using Wilcoxon paired t-test for matched pairs test or Kruskal-Wallis with Dunn’s multiple comparison test for comparing three or more groups. Significant *P* values were marked with stars, *P* <0.05 was represented by *, *P*<0.01 was represented by ** and *P*<0.001 was represented by ***.

## Results

### LeishIF4E2 is a cytoplasmic protein that contains an extended C-terminal region

Given the multiple eIF4E paralogs in *Leishmania*, and the assumption that they vary in their functional assignments, we examined the sequences of the different LeishIF4E paralogs, to evaluate their degree of homology and to search for non-conserved gaps and extensions. Figure S1 indicates that LeishIF4E2 indeed has a C-terminal extension, while LeishIF4E3 and LeishIF4E4 contain extended N-terminal regions (Figure S1). These N-terminal extensions have been widely examined for phosphorylation sites [19, 35, 36] and structural features [13]. The different LeishIF4Es vary from their mammalian counterpart, showing only around 29-40% similarity (Figure S1B). However, they also show large variations among themselves. The open reading frames of LeishIF4E2 from different *Leishmania* and *trypanosome* species were aligned using Jalview (2.10.5). The final alignment with predicted secondary structure was developed using the downloaded PDB file of the *Homo sapiens* IF4E1 (https://www.rcsb.org/) along with the generated FASTA alignment file, using the online ESPript 3 tool. The predicted structure of LeishIF4E2 is composed of alpha helices and beta strands. LeishIF4E2 contains an extended C-terminal region which is predicted to be highly disordered, in all *Leishmania* species. It is of interest to note that this extended C-terminal region is absent in trypanosome orthologs, in *T. brucei* and *T. congolensi* (Figure 1A).

**Fig 1.**
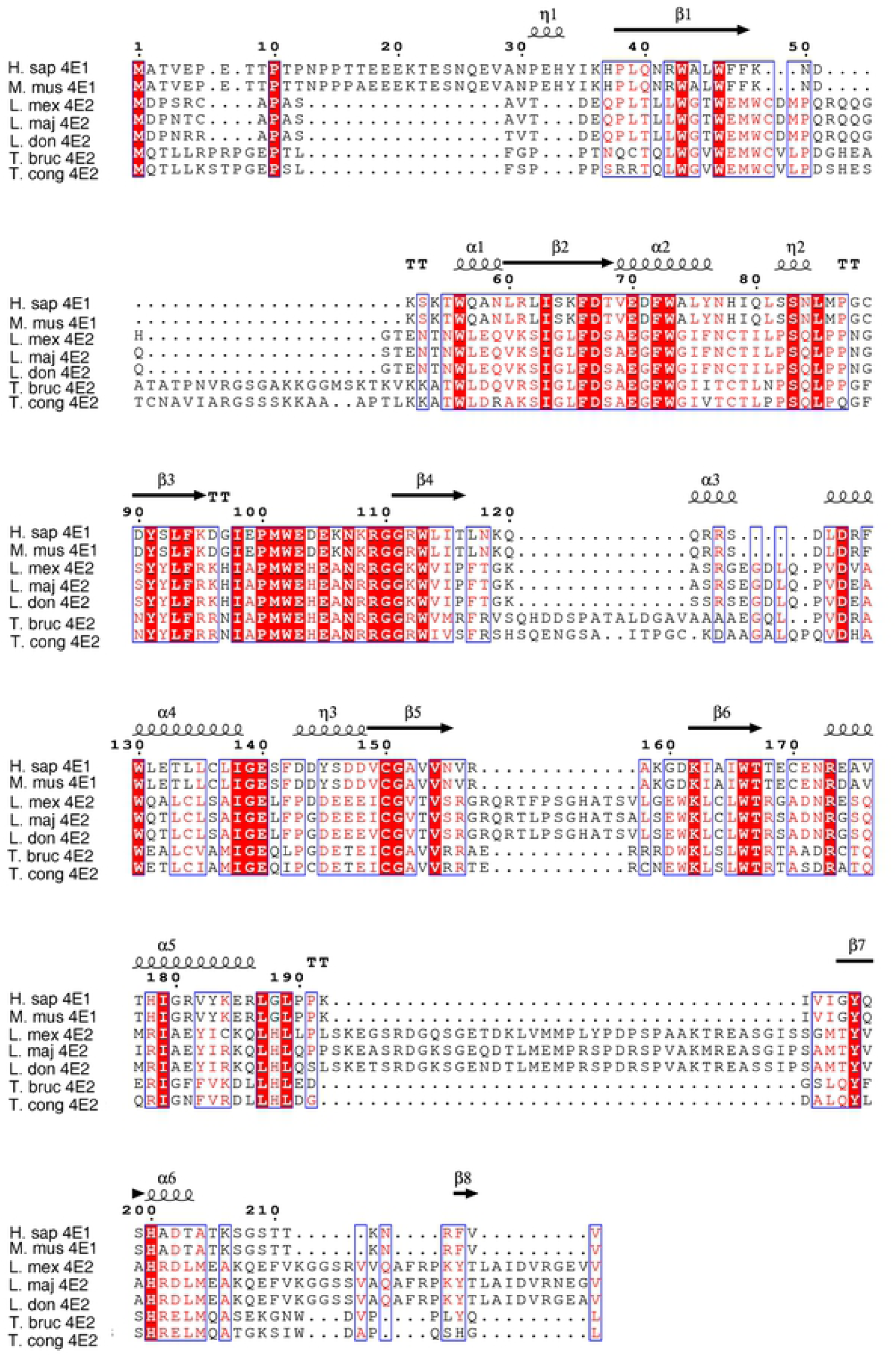
Multiple sequence alignment of the LeishIF4E2. The open reading frames of LeishIF4E2 from different *Leishmania* and *Trypanosoma* specis along with mammalian eIF4E1 were aligned using Jalview (2.10.5). The sequences were retrieved from *L. mexicana* (L. mex, LmxM.19.1480), *L. major* (L. maj, LmjF.19.1500); *L. donovani*, (L. don, LdBPK_191520.1), *T. brucei* (T. bruc, Tb927.10.16070) and *T. congolense* (T. con, TcIL3000_0_03820). The alignment file was saved in FASTA format. The final alignment showing the predicted secondary structure was developed using the downloaded PDB file (5EHC, DOI: <u>10.2210/pdb5EHC/pdb</u>) of *Homo sapiens* IF4E1 (https://www.rcsb.org/) along with the FASTA alignment file, using the online ESPript 3 tool [56].

It is noted that the major band that interacts with the specific antibodies against LeishIF4E2 migrates around 25 kDa, suggesting that it went through a cleavage process. To further investigate if the disordered region at the C-terminal end could lead to the potential cleavage product, full length recombinant LeishIF4E2, with an N-terminal Histidine tag was expressed in *E. coli* and affinity purified over a nickel column. The eluted fractions were examined by western analysis using an antibody against histidine. The affinity purified protein migrated as a major band of >25 kDa, and a minor higher band of <35 kDa. (Figure S1C). Since the protein was His-tagged at its N-terminus, based on the fragment size it appeared that the cleavage occurred just before the C-terminal domain.

We further investigated the localization of LeishIF4E2 within *Leishmania* cells. For this, SBP-tagged LeishIF4E2 was transfected into *Leishmania* cells and a stable cell line was selected by selection for drug resistance. The localization of LeishIF4E2 within the cells was determined using an immuno-histochemical approach, using confocal microscopy. Mid-log cells (1.2 x 10^7^ cells/mL) were harvested, washed and fixed in paraformaldehyde. The fixed cells were incubated with the primary anti-SBP antibody followed by a secondary anti-mouse antibody labeled with a fluorophore (Alexa Fluor 488 green). Nuclear and kinetoplast DNA were stained with DAPI. Figures 2 and S2 (Field view) show that, LeishIF4E2 is mostly and uniformly localized in the cytoplasm in an aggregative form.

**Fig 2.**
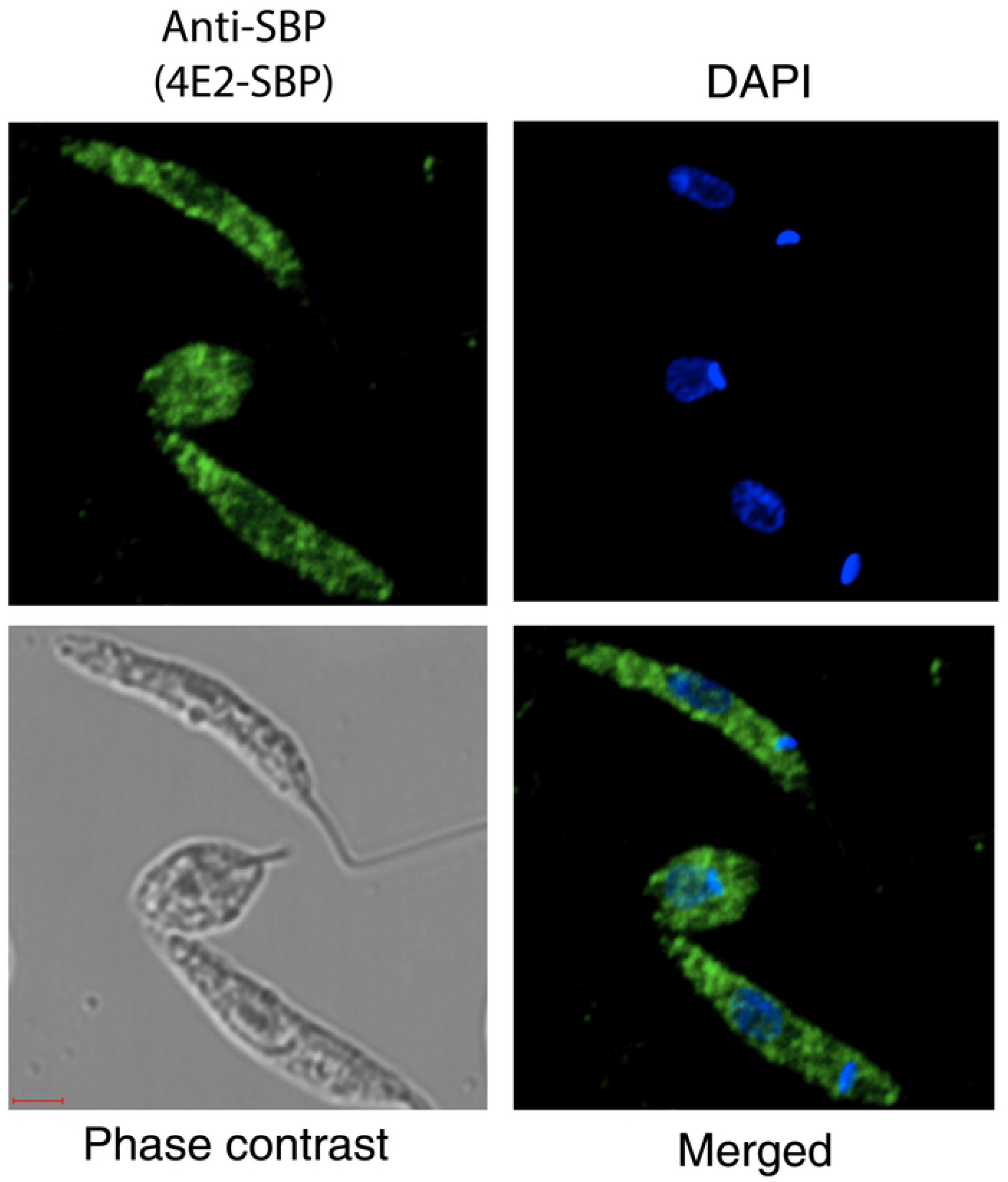
LeishIF4E2 is localized in the cytoplasm. *L. amazonensis* cells expressing LeishIF4E2-SBP were grown under normal conditions. The cells were washed, fixed in paraformaldehyde and further processed for confocal microscopy. LeishIF4E2 was detected using monoclonal the anti-SBP primary antibody and a secondary goat anti-mouse fluorescent antibody labeled with a green fluorophore (Alexafluore, 488 nM). The nuclear and kinetoplast DNA was stained using DAPI (blue). The images were merged. Images were taken using confocal microscopy showing a Z-projection that was produced by the Image J software. Scale bar: 2µm

### Deletion of a single copy of LeishIF4E2 by the CRISPR-Cas9

The tools used for functional genomic analyses in *Leishmania* received a strong boost by the development of the CRISPR-Cas9 system [25]. LeishIF4E2 is a less studied cap-binding protein paralog. We therefore attempted to delete its two alleles and examine how this deletion affected the overall physiology of the parasites. We first prepared a *L. mexicana* cell line that expressed Cas9 and T7 RNA polymerase, by transfection of the pTB007 plasmid (Beneke et al., 2017). The stably transfected cell line was selected for its hygromycin resistance. Specific sgRNAs targeted to LeishIF4E2 at the 5’ and 3’ UTRs that flanked the LeishIF4E2 ORF were generated by PCR, for cleavage around the target gene and its replacement with the G418 or Blastocidin repair fragments. The LeishIF4E2 deletion cell line was selected in the presence of G418 (200 µg/mL) and a diagnostic PCR analysis indicated that a single allele was eliminated. Our efforts to generate a null mutant were unsuccessful.

The diagnostic PCR was performed with genomic DNA of the mutant, using primers derived from the 5’ and 3’ UTRs of the LeishIF4E2 transcript. The reaction generated two products using gDNA of the mutant cells. These two products corresponded to a 1.2 Kb product that represented the endogenous LeishIF4E2 gene, and a second product of ∼2.1 Kb that corresponded to the integrated G418 resistance gene, flanked by the 5’ and 3’ LeishIF4E2 UTRs (Figure 3A left panel).

**Fig 3.**
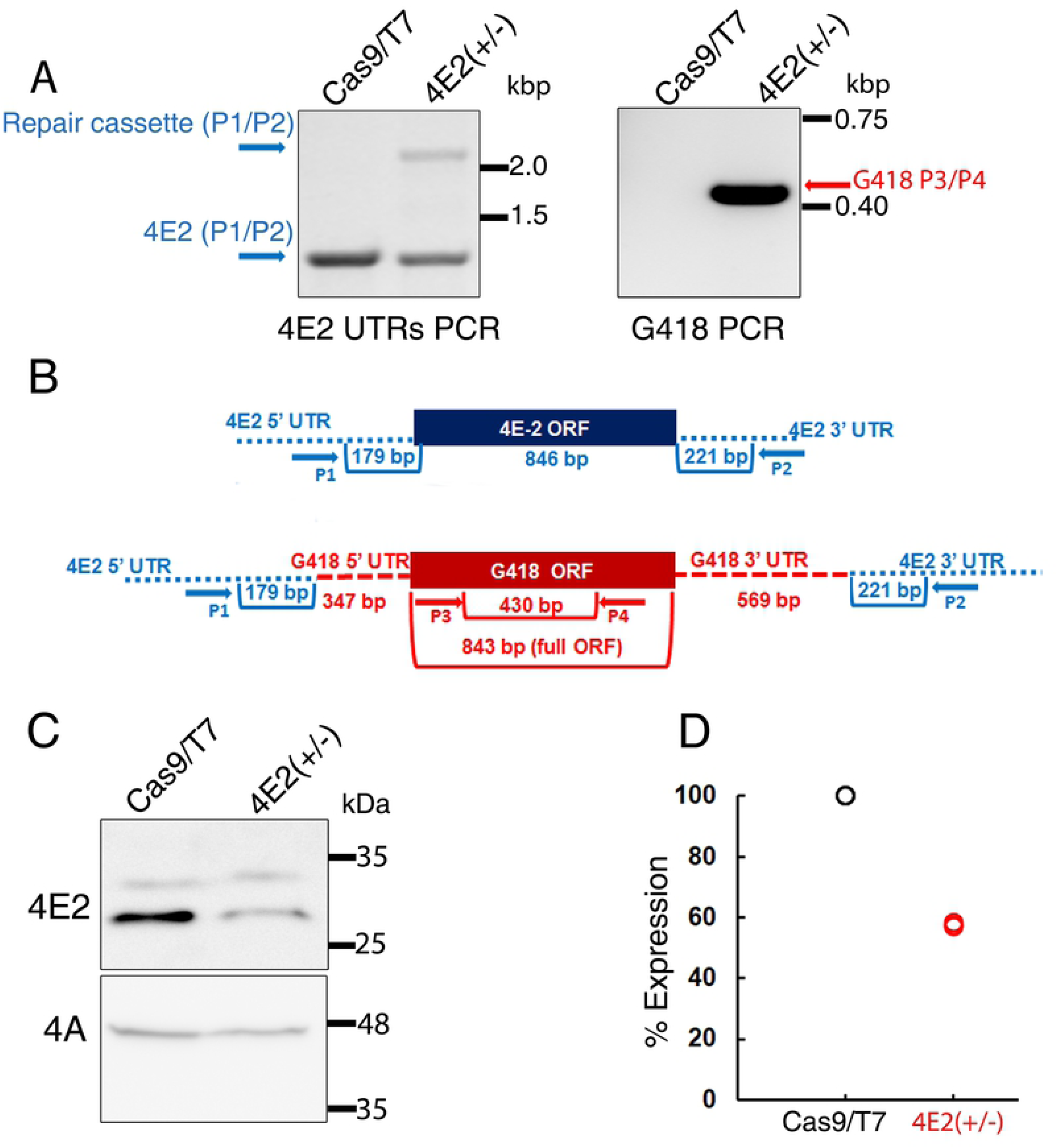
CRISPR-Cas9 mediated deletion of LeishIF4E2. **(A)** Diagnostic PCR was performed to confirm the deletion of single allele of LeishIF4E2. Genomic DNA of *L. mexicana* was extracted from the LeishIF4E2(+/-) mutant and from the *L. mexicana* Cas9/T7 cells. PCR was performed using primers derived from the LeishIF4E2 5’ and 3’ UTRs and from the ORF of G418 resistance gene. **(B)** Schematic representation of LeishIF4E2 locus and the primers (represented by arrows). The PCR was applied to test the presence or absence of the LeishIF4E2 gene and the G418 resistance marker. Primers derived from the LeishIF4E2 UTRs are shown in blue and primers derived from the ORF of G418 are shown in red. **(C)** Western analysis monitoring the protein level of LeishIF4E2 in the LeishIF4E2(+/-) mutant and in Cas9/T7 control was performed using LeishIF4E2 specific antibodies. LeishIF4A-1 served as a loading control. **(D)** Dot plot represents the densitometry analysis of the LeishIF4E2 protein levels in the LeishIF4E2(+/-) mutant and in the Cas9/T7 controls.

However, a similar PCR control reaction using the Cas9/T7 control gDNA as a template yielded only a single product (∼1.2 Kb), representing the endogenous LeishIF4E2 gene (Figure 3A left panel). We further verified the insertion of G418 repair fragment by using the G418 resistance gene specific primers which generated the expected product (450 bp) only with gDNA extracted from the LeishIF4E2(+/-) cell line, and not with Cas9/T7 control gDNA (Figure 3A right panel). Figure 3B represents the schematic design of the PCR primers used for the diagnosis purpose. These two PCR reactions confirmed the deletion of a single copy of the LeishIF4E2 gene from the *Leishmania* genome.

The effect of LeishIF4E2 heterozygous deletion on the target protein expression was examined by the western analysis of cell extracts obtained from the LeishIF4E2(+/-) mutant, as compared to control cells expressing Cas9/T7. The blots were reacted with LeishIF4E2 specific antibodies (Figure 3C upper panel). Antibodies against LeishIF4A served as a loading control to ensure equal loading (Figure 3C lower panel). Figure 3D shows a densitometry analysis of the LeishIF4E2 expression in the deletion mutant as compared to the Cas9/T7 control. LeishIF4E2 deletion mutant shows the 40% reduction in the protein levels as compared to control. We observed two bands in the western analysis, using antibodies specific for LeishIF4E2, one at ∼31 kDa and another above 25 kDa. A similar pattern of protein cleavage was observed with the recombinant LeishIF4E2(Figure S1C) and could originate from cleavage of the C-terminus. Mass spectrometry analysis correlated with the observed reduction in the protein level of LeishIF4E2 in LeishIF4E2(+/-) mutant cells, as compared to wild type (Table S1). The combination of the PCR, westerns and Mass spectrometry analyses validated the successful elimination of one copy LeishIF4E2 from the *L. mexicana* genome.

### The LeishIF4E2(+/-) deletion mutant shows altered promastigote morphology

Mid log phase (Day 2) Promastigotes of wild type, Cas9/T7 expressers, LeishIF4E2(+/-) cells, and LeishIF4E2 add-back cells grown at 25°C and neutral pH, were washed with PBS and fixed with 2% paraformaldehyde. The slides were visualized by phase contrast microscopy at 100x magnification. We noticed that the LeishIF4E2(+/-) deletion mutant possessed a defective morphology. The mutant cells became round, they reduced the length of their flagellum, and deviated from the typical promastigote form (Figure 4A). Control wild type and Cas9/T7 expressing cells exhibited normal promastigote features, comprising elongated shape that were equipped with a long and protruding flagella. Normal promastigote morphology was reverted back in LeishIF4E2(+/-) mutant when add-back SBP tagged-LeishIF4E2 was expressed by an episomal transfection with pT-Puro-H-LeishIF4E2-SBP-H (Figure S3). Once expression of LeishIF4E2 was recovered, cell morphology and flagellum growth were returned to their normal status, as observed in the wild type cells (Figure 4A right panel).

**Fig 4.**
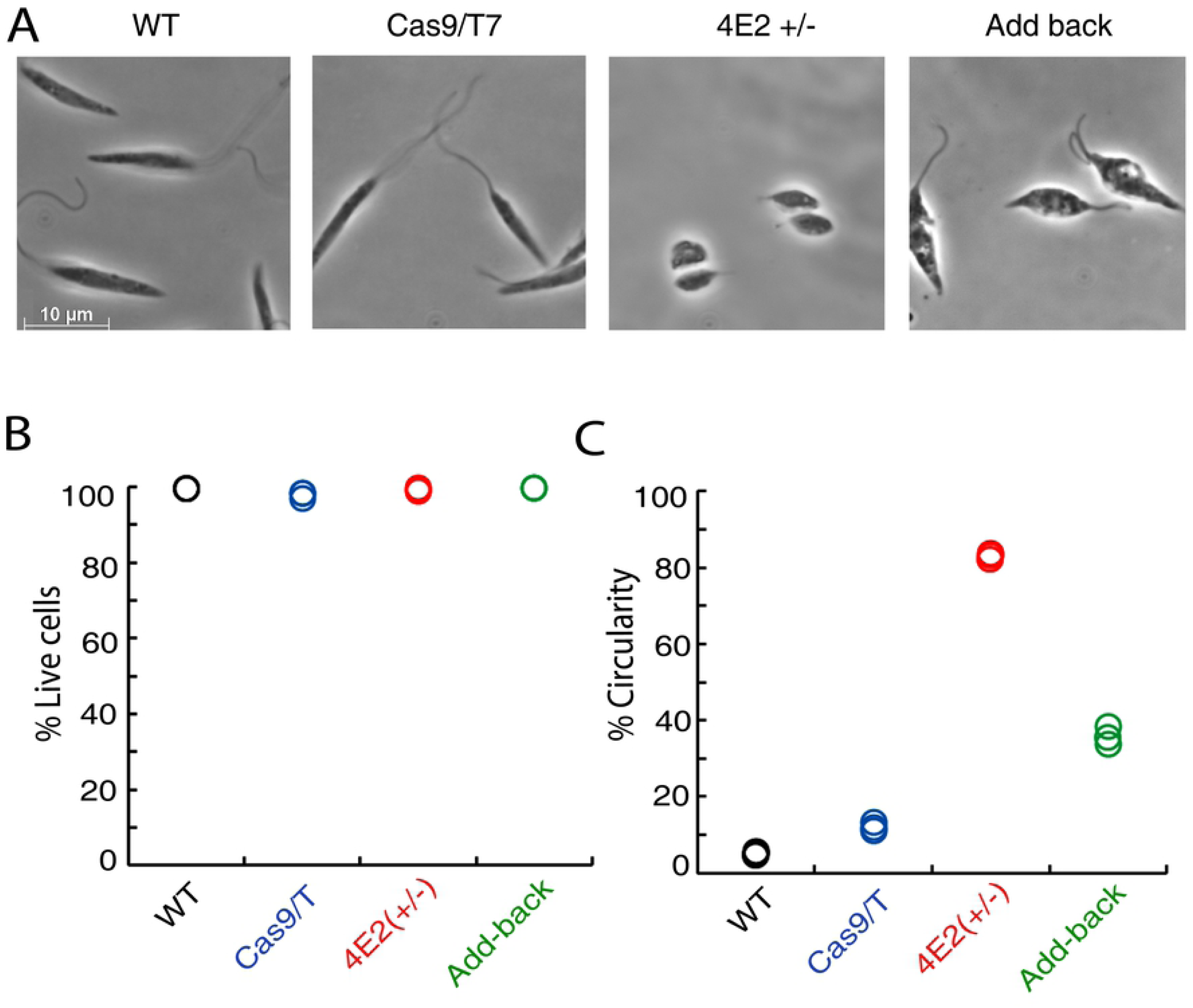
The LeishIF4E2(+/-) deletion mutant shows altered promastigote morphology. **(A)** Mid-log phase (Day 2) promastigotes of wild type, Cas9/T7 expressers, LeishIF4E2(+/-) cells, and LeishIF4E2 add-back cells were fixed with 2% paraformaldehyde and visualized by phase contrast microscopy at 100x magnification. Wild type (WT), Cas9/T7 expressing cells showed normal promastigote morphology while LeishIF4E2(+/-) became round with reduced flagellar length. This altered phenotype was reverted back to normal in LeishIF4E2 add-back cells. **(B)** All the cell lines were stained with 20 µg/mL propidium iodide (PI) for 30 minutes and cell viability was analyzed using the ImageStream X Mark II Imaging flow cytometer (Millipore). 20,000 cells were analyzed for each sample and percent viable cells were determined. **(C)**.The circularity of single, viable and focused cells from each of the cell lines was quantified using flow cytometry and shown as percentage of the total number of cells measured. Data from three independent experiments are shown.

We also used flow cytometry imaging to analyze the cell shape and viability of the cell populations. Cells were gated on the basis of being single (non-aggregated), focused (Figure S4B) and rounded up, and 20,000 cells from each group were analyzed. We measured cell viability by incubating with Propidium Iodide (PI) for 30 min. The viability of the LeishIF4E2(+/-) mutant was not affected as compared to control cells and cell viability was comparable between all the groups tested (Figure 4B and S4A). In parallel, we analyzed and quantified the changes in cell shape. In LeishIF4E2(+/-) mutant, around 83% cells were deviated from their normal promastigote morphology and they became round in shape with reduced flagella. In control wild type cells and Cas9/T7 expressing cells, only 5.14 % and 11.8% were circular shaped, respectively. The level of LeishIF4E2 was restored in the mutant cells, and in parallel we noticed a recovery from their mutant phenotype. The add-back LeishIF4E2-SBP show only 35% circular cells (Figure 4C and S4C).

### Global translation and growth are not affected by deletion of a single LeishIF4E2 allele

Since LeishIF4E2 is a cap-binding protein paralog, we investigated how the reduced level of LeishIF4E2 in the heterozygous mutant affected overall translation in the LeishIF4E2(+/-) mutant cells. We used the SUnSET assay to monitor general translation. The SUnSET assay is based on the incorporation of puromycin in the growing polypeptide chain, as it is a structural analogue of amino acyl tRNA that occupies the ribosomal A site. Integration of puromycin into the polypeptide chain blocks its elongation, resulting in translation termination. Control mid-log wild type, Cas9/T7 expressing cells, and LeishIF4E2(+/-) mutant cells were incubated with puromycin (1µg/mL) for 30 minutes. The cells were then harvested, extracted and resolved on SDS-PAGE. Puromycing incorporation was monitored by western analysis using anti-puromycin antibodies. A cycloheximide treated sample served as a negative control with no active puromycin incorporation.

Translation level in the LeishIF4E2(+/-) mutants was hardly affected by the allele deletion, as compared to control wild type and Cas9/T7 expressing cells (Figure 5A). Densitometry analysis of the western blot (Figure 5C), normalized to the total protein load (Figure 5B) showed that translation levels in Cas9/T7 and LeishIF4E2(+/-) mutants were 76.4% and 83.5%, respectively, as compared to wild type (100%). We further, monitored the growth rate of LeishIF4E2(+/-) mutants, as compared to the control cell lines. The *L. mexicana* LeishIF4E2(+/-) mutant, wild type and, Cas9/T7 expressing control cells, were cultured at 25°C in M199 containing essential nutrients, seeded at an initial concentration of 5×10^5^ cells/mL, and counted on a daily basis during four consecutive days. The growth curves (Figure 5D) show that the proliferation rate of the LeishIF4E2(+/-) mutant cells was comparable to that of the control wild type or Cas9/T7 expressing promastigotes. This is in agreement with the results obtained in the SUnSET assay, which demonstrates that general translation in the LeishIF4E2(+/-) mutant cells is not affected as compared to control cells.

**Fig 5.**
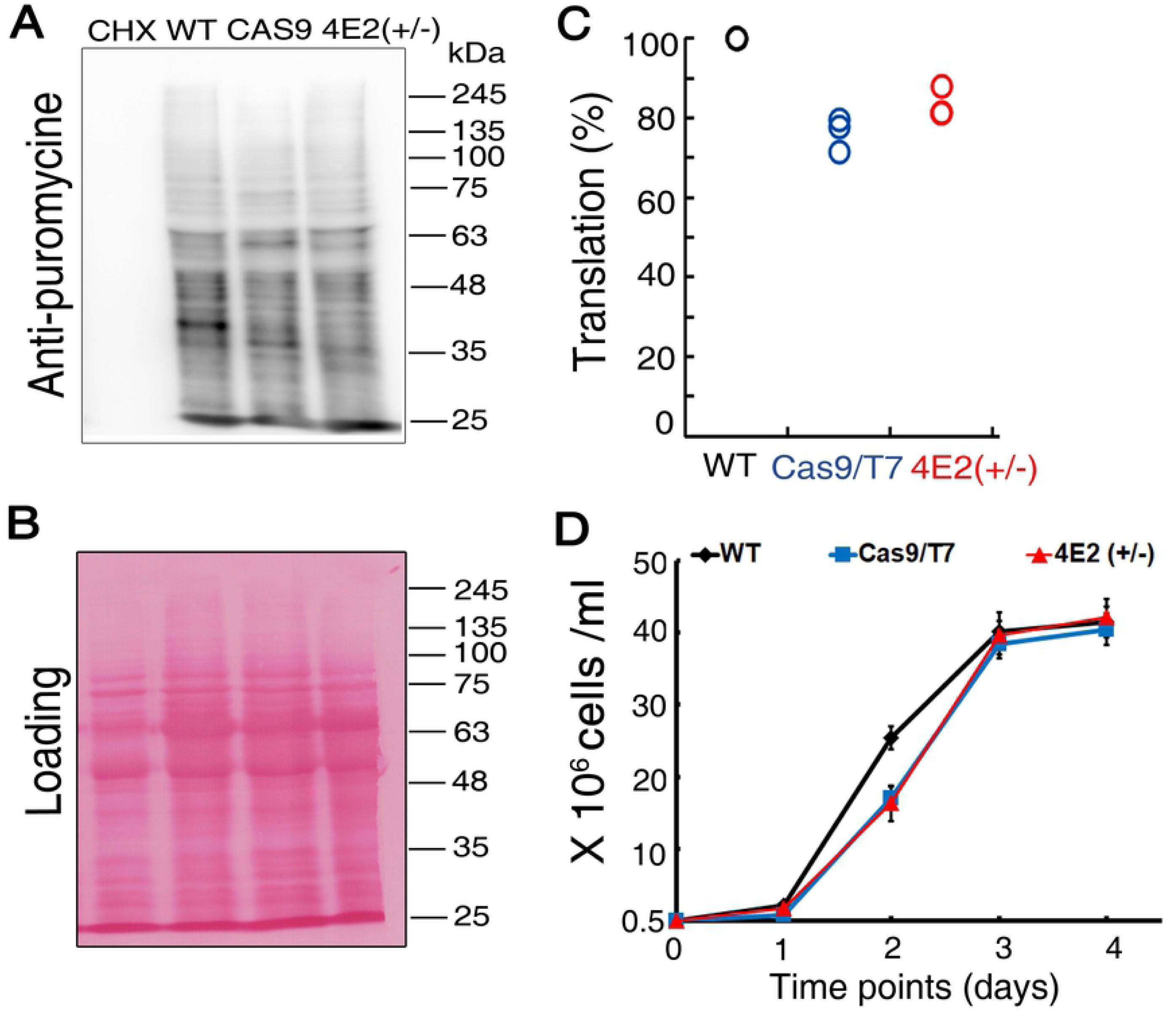
Global translation and growth curve is not altered in the LeishIF4E2(+/-) mutant cells. **(A)** LeishIF4E2(+/-) cells, wild type and Cas9/T7 expressing cells were incubated with 1 µg/mL puromycin for 30 min. Cycloheximide treated cells were used a negative control for complete inhibition of translation. Puromycin treated cells were lysed and resolved over 12% SDS-PAGE and subjected to western analysis using antibodies against puromycin. **(B)** Ponceau staining was used to indicate comparable protein loads. **(C)** Densitometry analysis of puromycin incorporation in the different cells lines, was compared to wild type (WT) cells (considered as 100%). Data from all three independent experiments are represented. WT cells are shown in black, Cas9/T7 is in blue and the LeishIF4E2(+/-) mutant cells are in red. **(D)** The *L. mexicana* LeishIF4E2(+/-) mutant, wild type, and Cas9/T7 expressing control cells, were cultured at 25°C in M199 containing essential nutrients. Cell counts were monitored daily during 4 consecutive days. The LeishIF4E2(+/-) mutant cells are shown in red, wild type cells in black, and Cas9/T7 expressing cells are shown in blue. The curves were obtained from three independent assays, error bars are also marked.

### The LeishIF4E2(+/-) mutant cells show reduced infectivity to macrophages

As the LeishIF4E2(+/-) mutant cells showed a defective morphology, due to their short flagellum and round body structure. We therefore examined their ability to infect cultured murine macrophage cells using the RAW 267.4 line. *Leishmania* parasites from respective cell lines were pre-stained with carboxyfluorescein succinimidyl ester (CFSE), and further used to infect the macrophages, at a multiplicity of 10:1 parasites per macrophage. Infection lasted for one hour, at 37°C. The macrophages were later washed to remove unbound parasites and the cells were fixed with paraformaldehyde (2%) and processed for confocal analysis. DAPI staining was used visualize the large macrophage nuclei. The infected macrophage cultures were examined in confocal microscopy either immediately (1 hr), or 24 hrs following infection. These two time points allowed us to keep track of initial entry of *Leishmania*, and further multiplication inside the host macrophages. Infectivity of the LeishIF4E2(+/-) mutant cells was compared to that of wild type cells and control Cas9/T7 expressing cells. We counted 100 macrophages from different fields, to determine the infectivity index. The results, shown in Figures 6A&B and S5A &B (for a broad field view) indicate that infectivity of the LeishIF4E2(+/-) mutant was impaired already 1 hr post infection as well as after 24 hrs. Infection by the LeishIF4E2(+/-) mutant was compared to that of wild type and Cas9/T7 expressers. Infection of the macrophages by wild type and Cas9-T7 parasites ranged between 88-99% of the cells. In both time points the infection of LeishIF4E2(+/-) mutant was significantly reduced as compared to control cells, showing 48% after 1 hr and 65% after 24 hrs (Figure S6 A&B). Infection by the add-back parasites showed a clear recovery in their ability to infect macrophages.

**Fig 6.**
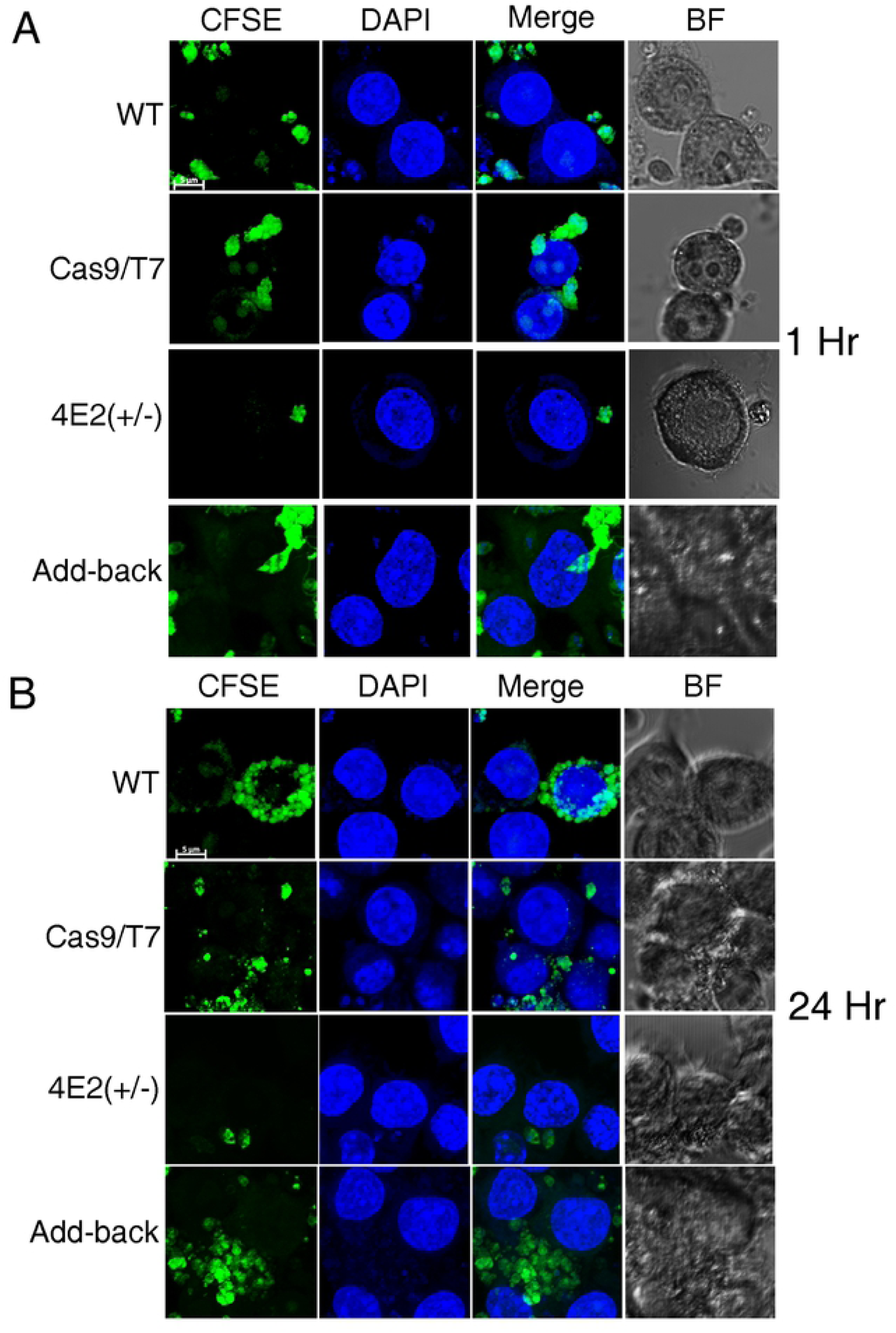
The LeishIF4E2(+/-) mutant cells shows reduced infectivity to macrophages. Stationary phase (Day 5) *L. mexicana* wild type (WT), Cas9/T7 expressing cells, LeishIF4E2(+/-) mutant cells and the add-back LeishIF4E2 line were pre-stained with CFSE (green), counted, washed and used to infected RAW 264.7 macrophages, at a ratio of 10:1 for one hour. The cells were then washed to remove unattached parasites, and the macrophages were cultured for 1h (A) or 24h (B) at 37°C. Macrophage nuclei were stained with DAPI and the infected macrophage slides were processed for confocal microscopy, showing a Z-projection produced by Image J software. Fields containing 100 cells were further evaluated to quantify the infection. The scale bar represents 5 µm.

We also counted the number of parasites per infected macrophage. A pronounced inhibitory effect of the LeishIF4E2(+/-) mutant was observed, due to a reduced number of parasites per infected macrophage (Figure 6 A&B, S5A&B, S6 A&B). While after 1 hr of infection wild type and Cas9/T7 control cells, each showed 2.5 and 2 parasites per infected macrophage, respectively, LeishIF4E2(+/-) cells showed only 1 parasite per infected macrophage at this time point. The reduced number of LeishIF4E2(+/-) parasites per infected cell was observed also at 24 hr post infection. Wild type and Cas9/T7 control cells showed 5.2 and 5 parasites per infected macrophage, respectively, and LeishIF4E2(+/-) mutant cells showed only 1.2 parasites per infected macrophage. The differences in the infectivity of the LeishIF4E2(+/-) mutant cells as reflected by both the reduced number of infected macrophages and in the reduced number of parasites per infected macrophage were significant when compared to infection by wild type, Cas9/T7 (S6 A&B).

### Proteomic analysis of the LeishIF4E2(+/-) mutant cells show upregulation of proteins involved in specific cellular processes and a reduction in cytoskeletal components as well as in ribosomal proteins

To examine potential differences in the proteomic profile of LeishIF4E2(+/-) mutant, we carried out mass spectrometry analysis of the total cell extracts in LeishIF4E2(+/-) mutant cells, as compared to wild type cells. Mid-log cells were used for this proteome analysis. We used three independent samples analyzed in parallel and in the same run. The *L. mexicana* genome in TritrypDB was used to assign the peptides to their source proteins, which were later quantified by the MaxQuant software. The proteomic content of the LeishIF4E2(+/-) cells was compared to that of control wild type cells, to identify proteins that were relatively increased or decreased, using a threshold of at least 1.6 fold, with p<0.05. Perseus software platform was used for the statistical analysis [[34], Table S1]. Figure 7A and Table S1 describe the manually categorized groups of 316 upregulated proteins and highlights a strong increase in the proteins that relate to various cellular processes. These included proteins categorized as involved in general cell metabolism, DNA replication and repair, signaling and finally cellular motor activity proteins. Metabolic proteins included mainly enzymes required for fatty acid and carbohydrate metabolism, while DNA replication and repair category contained mainly RAD50 repair enzymes. Various kinases and phosphatases were found within the signaling group. Finally, proteins related to the cellular motor activity group mostly included kinesins.

**Fig 7.**
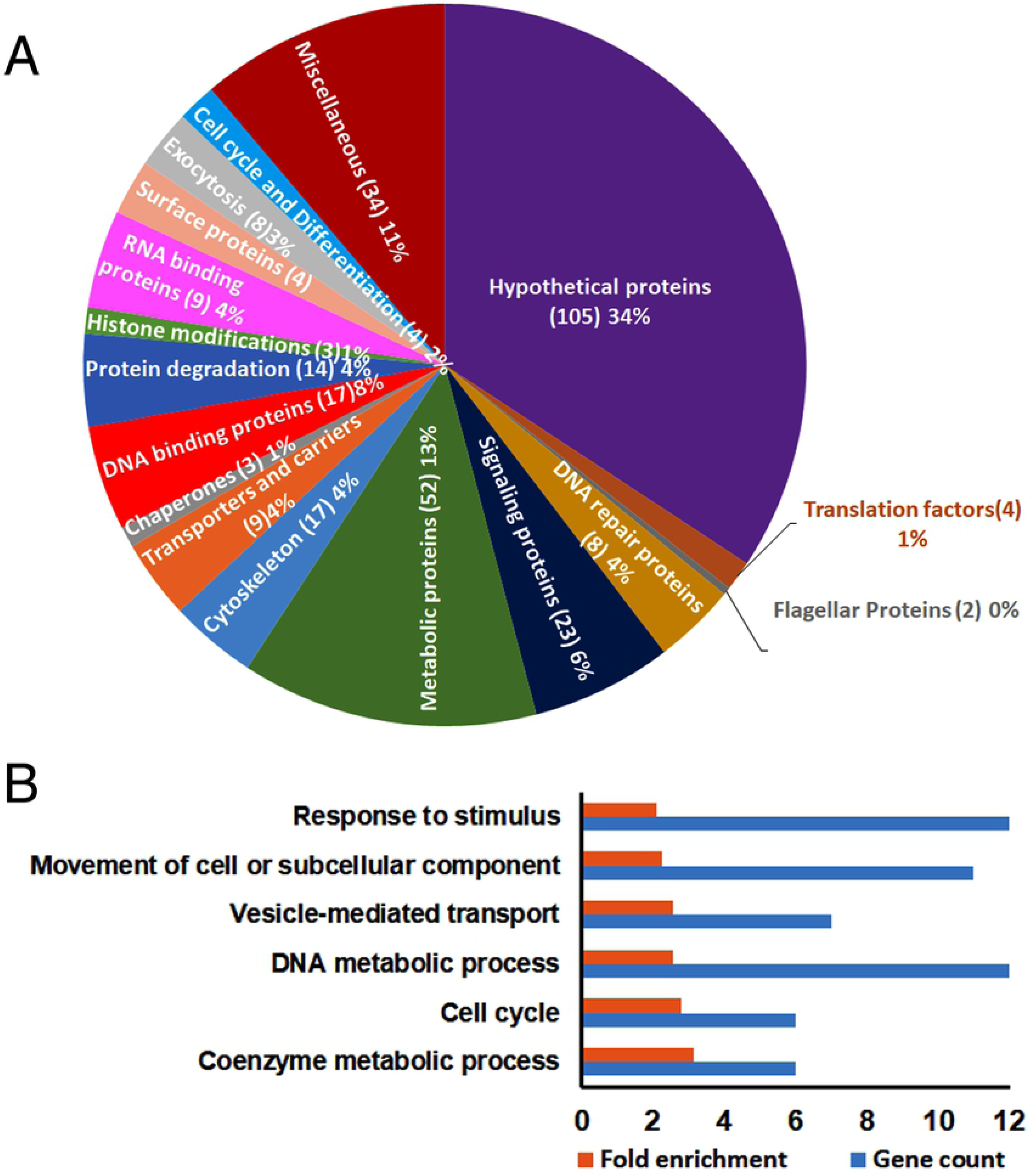
The categorized proteome of the upregulated proteins in LeishIF4E2(+/-) mutant cells as compared to WT. The proteomic content of LeishIF4E2(+/-) and WT cells was determined by LC-MS/MS analysis, in triplicates. Raw mass spectrometric data were analyzed and quantified using the MaxQuant software and the peptide data were searched against the annotated *L. mexicana* proteins listed in TriTrypDB. The summed intensities of the peptides that served to identify the individual proteins were used to quantify changes in the proteomic content of specific proteins. Statistical analysis was done using the Perseus software. Proteins that were upregulated in the LeishIF4E2(+/-) mutant by 1.6 fold as compared to WT cell extracts, with p<0.05 are shown. **(A**) Proteins in LeishIF4E2(+/-) that were upregulated (>1.6 fold) as compared to WT extracts were clustered manually into functional categories. The pie chart represents the summed intensities of upregulated proteins in the LeishIF4E2(+/-) mutant. Numbers in brackets indicate the number of proteins in each category. **(B)** Enriched proteins were classified by the GO enrichment tool in TriTrypDB, based on Biological Function. The threshold for the calculated enrichment of proteins based on their GO terms was set for 2.5 fold, with p<0.05. This threshold eliminated most of the general groups that represented parental GO terms. GO terms for which only a single protein was annotated were filtered out as well. In some cases, GO terms that were included in other functional terms are not shown, leaving only the representative GO term.

We also noticed a downregulation of 214 proteins in LeishIF4E2(+/-) mutant (Table S1). The downregulated proteins were mostly associated with the flagellum, the cytoskeleton and ribosomes. Other proteins that were reduced in LeishIF4E2(+/-) mutant included signaling and metabolic proteins. The relative level of surface proteins like GP46/PSA and hydrophilic acetylated surface proteins was also reduced. The latter is known to be involved in parasite virulence [37-40].

The upregulated proteins were also evaluated by the Gene Ontology (GO) enrichment analysis through the TryTripDB platform, based on their biological functions. Enrichment threshold was set at 2 fold. Figure 7B and Table S1 highlight the major categories of the upregulated protein groups, each containing at least 6 proteins. In line with the manually categorized proteins, the GO enrichment analysis also found that the upregulated proteins relate to the general cellular and DNA metabolism. Other groups that were upregulated in the GO enrichment analysis included the proteins involved in the cellular or subcellular movement.

### The LeishIF4E2(+/-) mutant shows normal differentiation to axenic amastigotes

The LeishIF4E2(+/-) mutant promastigotes looked small, round shaped and equipped with a very short flagellum (Figure 8A). This morphology is similar to that observed during differentiation to axenic-like cells. We further examined how the mutant cells responded to conditions known to generate axenic amastigotes. L. *mexicana* promastigotes from wild type, Cas9/T7 expressing cells and LeishIF4E2(+/-) mutant were grown to reach their late log phase, and then transferred to conditions that induce differentiation to axenic amastigotes *in vitro* (pH 5.5, 33°C). These cultures were allowed to differentiate during four days. The differentiated LeishIF4E2(+/-) cells maintained a similar morphology, except that they became smaller, resembling axenic amastigote structure (Figure 8B).

**Fig 8.**
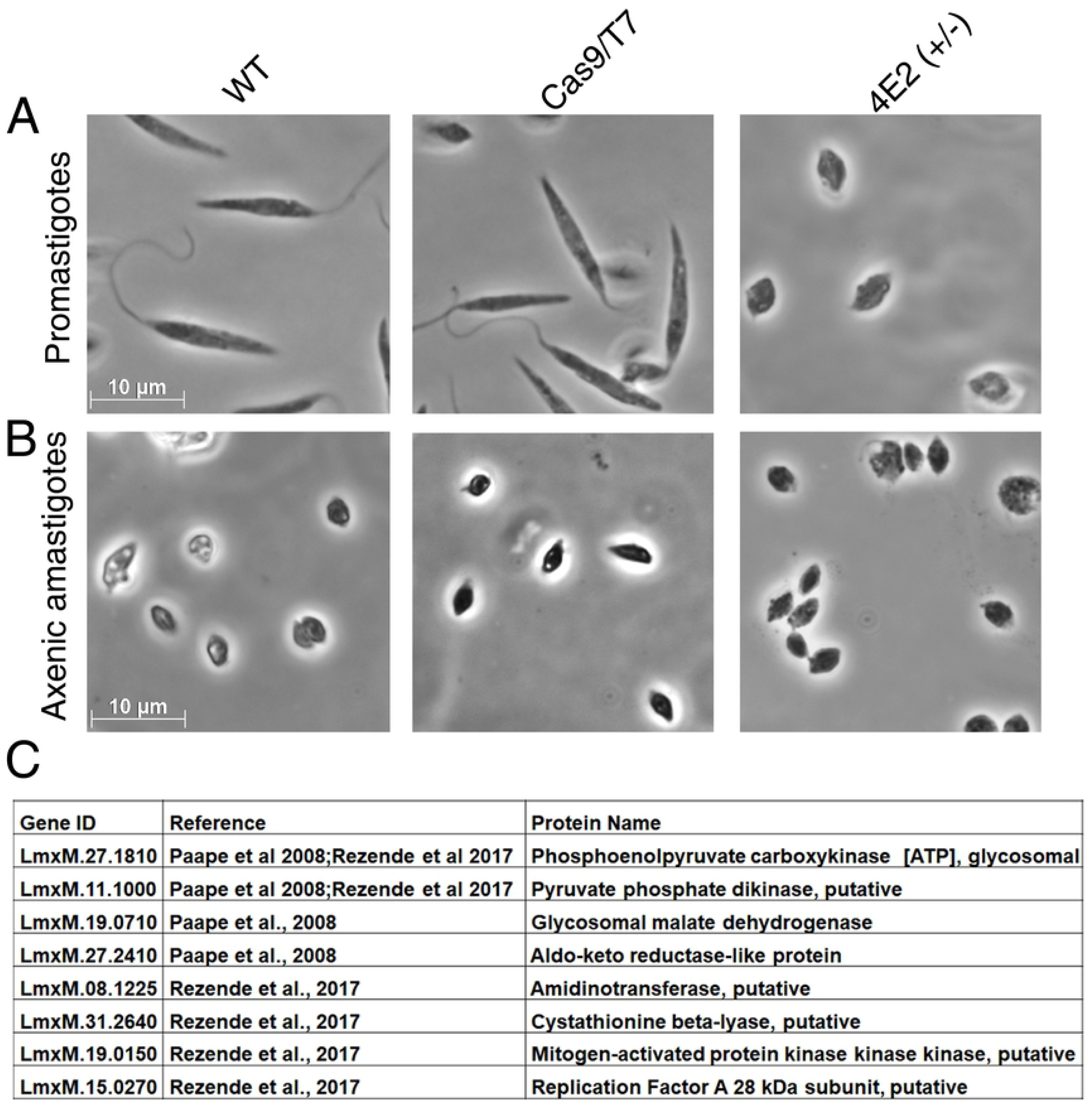
LeishIF4E2(+/-) mutant cells easily transform to axenic amastigote-like cells. **(A)** Promastigotes of the LeishIF4E2(+/-) mutant, control wild type, and Cas9/T7 expressing cells grown under normal conditions are shown. **(B)** Morphology of cells transferred to conditions that induce differentiation to axenic-amastigotes (33°C/pH 5.5) during four days. Images were captured at100x magnification with a Zeiss Axiovert 200M microscope equipped with AxioCam HRm CCD camera. The scale bar is 10 µm. **(C)** Upregulated proteins in the mutant LeishIF4E2(+/-) promastigotes (compared to WT cells) and in published amastigotes proteome. The total protein of the LeishIF4E2(+/-) mutant promastigotes was compared to the proteome of WT cells. The list of upregulated proteins was further compared with the proteins enriched in the amastigote proteome of virulent *L. amazonensis* PH8 strain (de Rezende et al, PLOS NTD 2017). and of *L. mexicana* amastigotes (Paape et al, Mol Cell Prot 2008).

We compared the repertoire of proteins that were enriched in the LeishIF4E2(+/-) mutant promastigotes with the published proteome that is specific to amastigotes (Paape etal 2008; Rezende et al 2017). This comparison highlighted only eight proteins that were upregulated in LeishIF4E2(+/-) mutant and also found in the amastigote specific proteins (Figure 8C and Table S2) that were reported previously [41, 42]. The shared proteins included important enzymes required for gluconeogenesis, such as the phosphoenolpyruvate carboxykinase and pyruvate phosphate dikinase. These two enzymes are key players in the gluconeogenesis pathways, which is an active pathway in the amastigote stage of *Leishmania*, and are required for the intracellular survival of the parasite within mammalian host cells (Rodriguez-Contreras et al., 2014). We also noticed that the aldo-keto reductase was upregulated in LeishIF4E2(+/-). Aldo-keto reductase is suggested to have a role in the removal of the ketoaldehyde metabolites derived from lipids, trioses (Roberts et al., 2018). Other proteins upregulated in LeishIF4E2(+/-) includes cystathionine beta-lyase (Methionine biosynthesis) and signaling kinases like mitogen-activated protein kinase. Enzymes involved in the lipid metabolism pathways were shown to increase in *Leishmania* amastigotes, as these change their energy source when they enter into mammalian macrophages [43].

## Discussion

The current study reports on our attempt to understand one of the least studied *Leishmania* cap-binding paralogs, LeishIF4E2, using the CRISPR-Cas9 mediated gene silencing methodology. We successfully deleted one of the two LeishIF4E2 alleles and studied the effect of this deletion on various cellular processes, including growth, morphology metabolism, infectivity to macrophages and overall proteome.

LeishIF4E2 does not associate with any eIF4G partner, and co-migrates with polysomes on sucrose gradients [10]. This profile does not lead to a definite conclusion on its role. On one hand, it reminds of the translation repressor 4E-HP [44], but on the other hand, we recently showed that LeishIF4E1 does not appear to function as a translation repressor, despite its inability to partner with an eIF4G candidate [15]. However, the possibility of having a dual role in translation, as a putative repressor or inducer for specific transcripts is considered as well.

The ortholog of LeishIF4E2 from *T. brucei*, TbEIF4E2, was shown to associate with a stem-loop binding protein that binds to histone mRNA, but its role in translation activation or repression remained elusive. Our current study uses a LeishIF4E2 mutant to explore the potential role of LeishIF4E2 during translation in *Leishmania*. Using the CRISPR-Cas9 technique for gene knockout, we generated a heterozygous mutant of LeishIF4E2, and examined how its deletion affected the mutant proteomic profile. Attempts to delete the two copies of LeishIF4E2 were unsuccessful, despite numerous efforts. This could be due to technical difficulties but we also cannot exclude the possibility that LeishIF4E2 is essential for survival. This is further supported by the fact that we were able to generate a null mutant of another cap-binding protein, LeishIF4E1, using this same approach [15]. The deletion of a single copy of LeishIF4E2 did not affect the general translation and growth rates in the mutant cells. However, the absence of one gene copy led to a change in the profile of the cell proteome. We noticed the upregulation of 316 proteins in LeishIF4E2(+/-) cells, while only 214 proteins were downregulated in this mutant, as compared to the wild type control. The GO enrichment analysis demonstrated that the groups showing decreased expression were categorized as flagellar and cytoskeletal proteins, in accordance with the accumulated defects in its morphology.

The LeishIF4E2 sequences are highly conserved among the *Leishmania* species, and less with trypanosomes. LeishIF4E2 possesses an extended C-terminal domain, which is predicted to be disordered. This extended C-terminus is absent from all other LeishIF4Es, emphasizing this feature as unique to LeishIF4E2. It was further noted that the extended C-terminus is also absent from the *T. brucei* and *T. congolensi* orthologs of LeishIF4E2. Thus, this C-terminus is specific to the *Leishmania* LeishIF4E2. We also show that the recombinant LeishIF4E2 purified from bacterial extracts is subject to proteolyic cleavage, generating two products, one of them lacking the C-terminal domain, as confirmed by the interaction with antibodies against the N-terminal His-tag. However, the biological role of this cleavage and its products needs to be further investigated.

It is of interest that complete or partial deletion of other LeishIF4E paralogs, resulted in a similar effect on cell morphology, as the mutant cells were small, non-flagellated and impaired in their ability to infect macrophages. This was the case with the null mutant of LeishIF4E1[15] and the heterozygous mutant of LeishIF4E3 [45]. Currently, the heterozygous deletion of LeishIF4E2 also generates morphologically defective cells (this study). In all these cases, we observed a similar pattern of behavior, namely that perturbation of individual LeishIF4Es, resulted in altered morphology, defects in flagellar growth and impaired infectivity. Based on these observations we can suggest that the different LeishIF4Es could be involved in control of the cytoskeleton and flagellar growth. It is also expected that the flagellum is required for the entry of promastigotes into macrophages once transmitted into the mammalian host, based on previous reports that the flagellum is involved in the parasite infectivity [46-48]. Apart from the defective morphology, the downregulation of important virulent factors such as discrete surface proteins, promastigote surface antigens (PSA/GP46), could result in the impaired infectivity. Expression of the GP46/PSA surface antigen that was previously shown to be involved in the parasite virulence by protecting the parasite from complement mediated lysis [40], was also reduced.

Genome wide tethering screens for identifying regulatory proteins that affect gene expression in *T. brucei* suggested that TbEIF4E2 is a translation repressor [49]. Our results are in line with this conclusion, since the translation assays monitoring de novo translation in the LeishIF4E2(+/-) mutant cells, did not show any decrease in protein synthesis activity, despite the reduced expression of LeishIF4E2 in this mutant. Furthermore, analysis of the LeishIF4E2(+/-) mutant proteome indicated an upregulation of 316 polypeptides, whereas only 214 proteins were downregulated. Given these results, we are considering different possibilities. One is that LeishIF4E2 could function as a translation repressor, as indicated above in *T. brucei.* Alternatively, LeishIF4E2 could function as a transcript specific translation factor, explaining why its absence was not reflected in the general translation profile. We also cannot exclude the option that the reduced expression of LeishIF4E2 had an indirect effect on the proteomic profile, through modulation of a small number of proteins involved in translation. Opposite roles for a specific cap-binding protein was reported for the 4E-HP ortholog in higher eukaryotes. This protein fails to bind any eIF4G partner, and was reported to function as a translation repressor during embryogenesis [50]. However, it was also reported to function as translation inducing factor for a specific set of transcripts that are translation under hypoxia conditions [51].

We further considered the possibility that LeishIF4E2 is associated with stage differentiation. Our former publication [14] indicates that LeishIF4E2 expression is almost undetectable in axenic amastigotes. Other cap-binding proteins vary, as LeishIF4E1 maintains its expression level at both life stages, while LeishIF4E-4 changes its modification profile, and LeishIF4E-3 is reduced in axenic amastigotes. The complete absence of LeishIF4E2 expression in axenic amastigotes could indicate that stage transformation requires its suppression during differentiation. We now see that reduction of LeishIF4E2(+/-) expression leads to upregulation of important enzymes of gluconeogenesis, as compared to wild type cells. This group of proteins was also reported to increase in amastigotes [52, 53]. Another protein that upregulates when LeishIF4E2 decreases its expression is the mitogen-activated protein kinase-kinase (LmxM.19.0150), a protein that was reported to increase in axenic amastigotes in previous publications [53]. The *T. brucei* ortholog, TbEIF4E2 was shown to bind the stem-loop binding protein 2, SLBP2. This protein associates specifically with histone mRNAs. It is of interest to note that in higher eukaryotes SLBP associates with polyribosomes, as a result of continued synthesis and transport of the histone mRNP to the cytoplasm [54]. In accordance, LeishIF4E2 is the only cap-binding protein that clearly associated with polysomal fractions that were separated over sucrose gradients, as other LeishIF4Es tested, co-migrated with higher fractions [10]. The repertoire of upregulated genes in the LeishIF4E2 (+/-) mutant shows enrichment of DNA-repair and DNA-binding proteins, which are also involved in DNA replication. These proteins were not upregulated in the LeishIF4E1(-/-) null mutant cells [15], emphasizing perhaps unique roles for the different cap-binding proteins.

The results shown in this study, and compared to our previous papers that described the effects of eliminating expression of specific cap-binding proteins and to studies in *T. brucei*, further emphasize the complex nature of the network that regulates translation in trypanosomatids. Results that are partially conclusive may also indicate on overlapping functions of the different cap-binding proteins. One should also consider the involvement of 4E-interacting proteins, such as Leish4E-IP1 [14] and Leish4E-IP2 [55] in modulating protein synthesis in this group of fascinating organisms that diverged early during evolution of eukaryotes.

## Legends to supplemental materials

**S1 Fig. A. Sequence alignment of the cap-binding protein paralogs of *Leishmania mexicana* indicating the N- and C-termini extensions.** Sequence alignment was generated using Jalview (2.10.5). The aligned sequences were derived from *L.mexicana* genome sequences from TritrypDB. The sequences used were LeishIF4E1 (4E1, <u>LmxM.27.1620</u>); LeishIF4E2, (4E2, LmxM.19.1480); LeishIF4E3 (4E3, LmxM.28.2500); LeishIF4E4 (4E4, <u>LmxM.29.0450</u>); LeishIF4E5 (4E5, LmxM.36.0590); LeishIF4E6 (4E6, <u>LmxM.26.0240</u>). N-terminal extensions are observed in LeishiF4E-3 and LeishIF4E-4, as previously reported. A C-terminal extension is observed in LeishIF4E-2. **B. Comparison of LeishIF4Es with mouse counterpart**-Table showing % similarity of LeishIF4 paralogs to mouse eIF4E. **C. Recombinant LeishIF4E-2**. Full length Histidine tagged LeishIF4E-2 at N-terminal end was expressed in BL-21 cells, purified with Nickel column and resolved over SDS-PAGE and subjected to western analysis using specific antibodies against Histidine.

**S2 Fig. LeishIF4E-2 is localized in the cytoplasm (Field)**. *L. amazonensis* cells expressing LeishIF4E2-SBP were grown under normal conditions. The cells were washed, fixed in paraformaldehyde and further processed for confocal microscopy. LeishIF4E-2 was detected using monoclonal anti -SBP primary antibody and a secondary goat anti mouse fluorescent antibody labeled with a green fluorophore (Alexafluore, 488 nM). The nuclear and kinetoplast DNA was stained using DAPI (blue). Images were taken using confocal microscopy showing a Z-projection that was produced by the Image J software. Scale bar: 10µm

**S3 Fig. Confirmation of add-back LeishIF4E2 expression. (A)** Cell lysates of *L. mexicana* LeishIF4E2 add-back and wild type cells were resolved over 10% SDS-PAGE followed by western analysis with antibodies directed against SBP (4E1). **(B)** The Ponceau staining of Tubulin on the blots served as a loading control.

**S4 Fig. Flow cytometry for viability, gating of focused single cell population and cell shape quantification.** *L. mexicana* wild type, Cas9/T7 expressing control cells, LeishIF4E-2(+/-) deletion mutant and add-back promastigotes were subjected to Flow cytometry analysis. **(A)** Cell viability is represented for focused, single gated cells for all the different cell lines **(B)** Scatter plots representing gated focused single cell populations for different cell lines. **(C)** Cell shapes are being represented in terms of circularity or elongatedness as scatter plots for gated cell population.

**S5 Fig. The LeishIF4E-2(+/-) mutant cells shows reduced infectivity to cultured macrophages (field).** Stationary phase *L. mexicana* LeishIF4E-2(+/-) mutant, wild type and Cas9/T7 expressers along with add-back cells, were pre-stained with the CFSE dye and further used to infect RAW 264.7 macrophages, at a ratio of 10:1 for one hour. The cells were then washed to remove excess parasites, and the macrophages were cultured for 1 hr (A) or 24 hr (B) post infection at 37°C. Macrophage nuclei were stained with DAPI and the infected macrophages were processed for confocal microscopy. A representative section of Z-projections (maximum intensity) produced by Image J software is presented in the figure. Fields containing 200 cells were further evaluated to quantify the infection.

**S6 Fig. Statistical analysis of the LeishIF4E-2(+/-) mutant infectivity as compared to controls**. Parasite infectivity of cultured RAW 264.7 macrophages was estimated *in vitro*, using the Image J software. **(A)** The percentage of infected macrophages was determined by counting a total of 100 macrophages from three independent experiments**. (B)** The average number of parasites per infected cell is shown. Kruskal Wallis test in GraphPad Prism was performed for the statistical analysis of the percentage of infected cells and for calculation of the average parasites per cell, along with standard deviation values (SD). The percentage of infected macrophages (%) and the average number of parasites per macrophage in the LeishIF4E-1(-/-) deletion mutant were compared with each of the control lines: wild type, Cas9/T7 expressing cells and LeishIF4E-1 add-back. *P* value < 0.001 is represented by *** while P value < 0.01 is represented by ** and P value < 0.05 is represented by *. The statistical difference between the control lines were non-significant. The data is shown for 1hr and 24 hr macrophage infections in separate panels.

**S1 Table. List of proteins identified in the total extract of 4E2(+/-) mutant and wild type *Leishmania mexicana.*** Raw mass spectrometric data were analyzed and quantified using the MaxQuant software and the peptide data were searched against the annotated *L. mexicana* proteins listed in TriTrypDB. The proteomic content of 4E2(+/-) and WT cells was determined by LC-MS/MS analysis, in triplicates

**S2 Table. Comparison of the 4E2(+/-) proteome with published amastigote proteomes.** The proteome of upregulated 4E2(+/-) mutant promastigotes was compared with the proteins enriched in the amastigote proteome of virulent *L. amazonensis* PH8 strain, as compared to the less virulent LV79 [57].

## Authors’ contributions

NT, NB and MS conceived the study. NT and NB contributed equally. MS is the principal investigator and RZ is the co-investigator. Both reviewed and edited the manuscript and participated in securing funding. NT and NB designed and performed the experiments with guidance of MS and wrote the manuscript. GD was involved in the cloning and purification of LeishIF4E-2. The funders had no role in study design, data collection and interpretation, or the decision to submit the work for publication

## Acknowledgments

We thank Dr. Eva Gluenz from the University of Oxford for providing us with the plasmids for performing CRISPR-Cas9 in *Leishmania.* We thank Dr. Uzi Hadad from the Ilse Katz Institute for Nanoscale Science & Technology at the Ben-Gurion University of the Negev for help with flow cytometry analysis and confocal microscopy. We thank Prof. Charles Jaffe (Hebrew University, Israel) for providing us with the *L. mexicana* strain M379. We thank the Smoler Proteomics Center in the Technion, Haifa, for their professional proteomic analysis.

